# Forecasting virus diffusion with simple Bayesian forecast combination

**DOI:** 10.1101/2020.06.10.144006

**Authors:** Philip Hans Franses

## Abstract

There are various diffusion models for S shaped processes like virus diffusion and these models are typically not nested. In this note it is proposed to combine the forecasts using a simple Bayesian forecast combination algorithm. An illustration to daily data on cumulative Covid-19 cases in the Netherlands shows the ease of use of the algorithm and the accuracy of the thus combined forecasts.

## 1. Introduction

Virus diffusion often obeys an S shaped pattern. Consider for example the daily cumulative new cases of Covid-19 in the Netherlands in Figure 1, for the sample February 27, 2020 with the first case, and May 19, 2020 with 44249 cases. The S shape is obvious. Figure 2 presents the daily new cases, and, as expected, a humped shape is noticeable, although at the same time variation across the days can be large.

**Figure 1:**
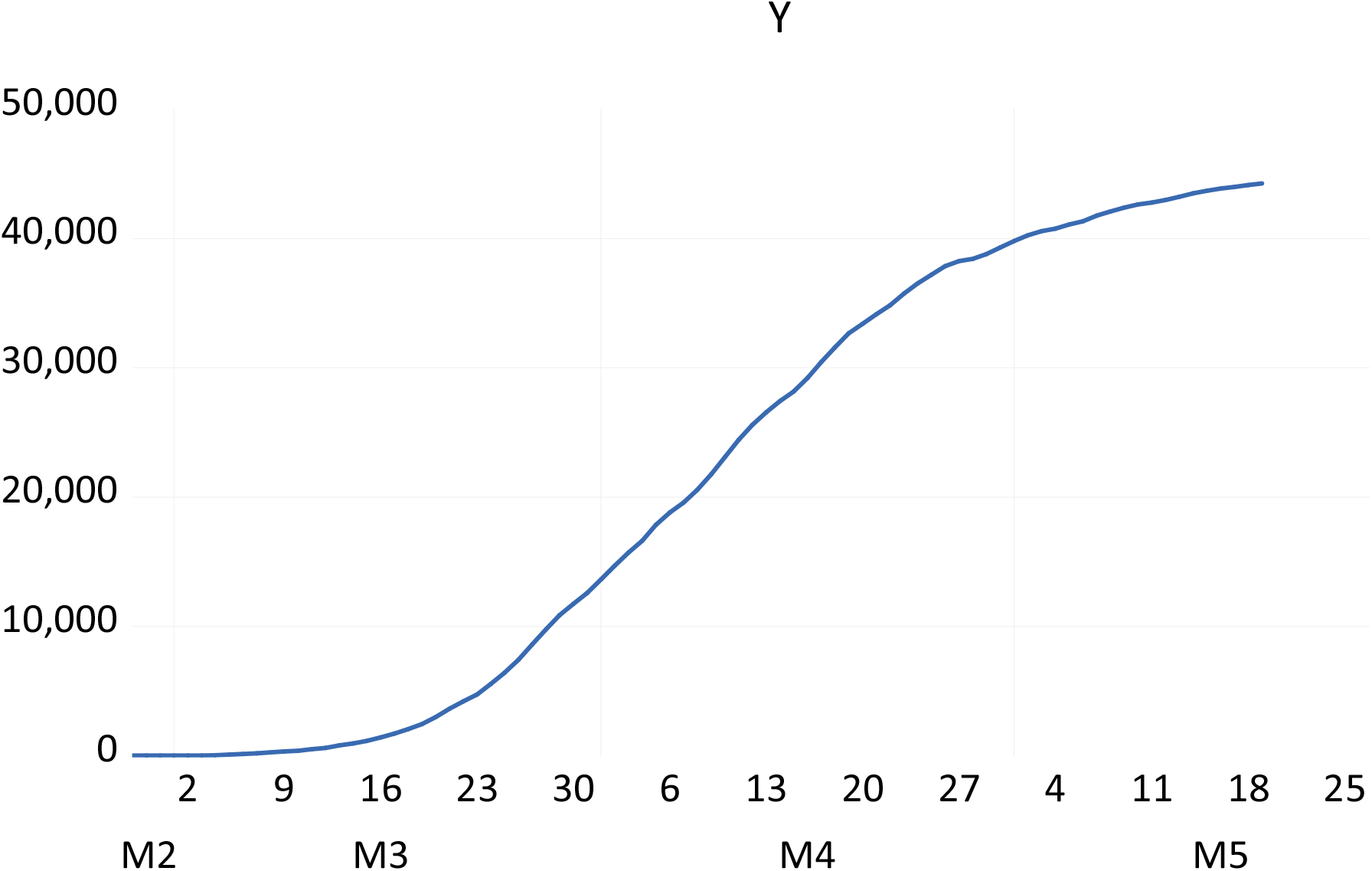
Cumulative new cases with Corona in the Netherlands until and including May 19, 2020, where February 27, 2020 marks case 1.

**Figure 2:**
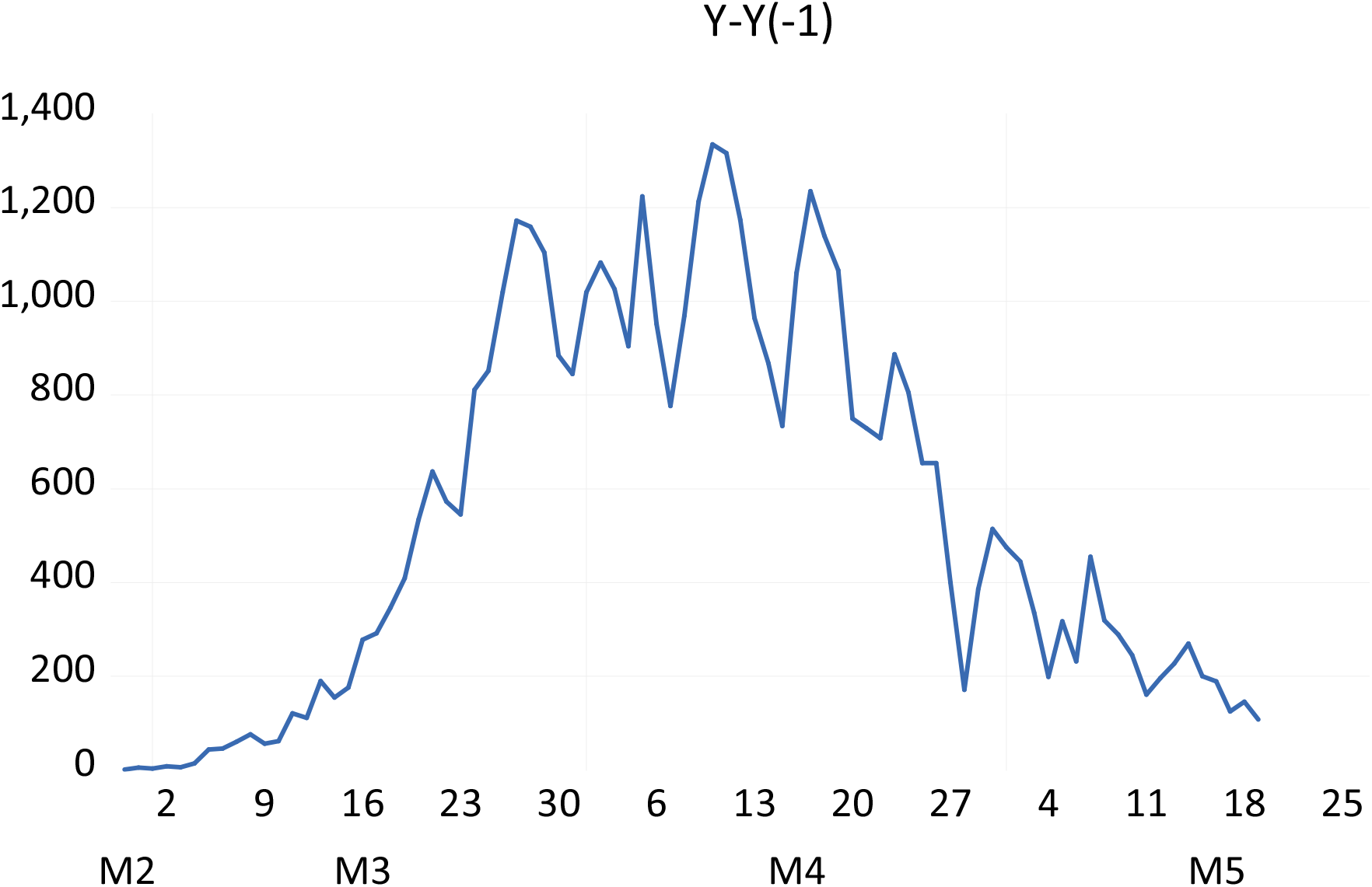
Daily new cases with Corona in the Netherlands, until and including May 19, 2020, where February 27, 2020 marks case 1.

Each day it can be of interest to predict the total cumulative number of cases for the next day. There are various models for S shaped data, and there is no evidence that one of these models is always preferable over the others. Hence, it seems reasonable to consider various models at the same time. For forecasting, it may then also be useful to combine the forecast, see Bates and Granger (1969), Clemen (1989) and Timmermann (2006) for evidence that combined forecasts often yield an improvement over individual forecasts. A typical combination refers to assign equal weights to the forecasts. In this paper it is however proposed to assign weights based on in-sample performance of each of the models, as reflected by their posterior probabilities, see Leamer (1978) for the first discussion of Bayesian model averaging. In practice, Bayesian methods may sometimes be difficult to handle, but here I rely on an important result in Raftery (1995, page 145), which is that the posterior probability can be expressed as a simple function of the values of the Bayesian Information Criterion (BIC) (Schwarz 1978). Such a BIC value is easily with the estimated maximum likelihood and many statistical packages report BIC values.

The outline of this note is as follows. Section 2 present three models that are frequently used for modeling and forecasting S shaped diffusion processes. Section 3 presents the simple algorithm to compute combined forecasts. Section 4 analyzes the recursive one-day-ahead forecasts for the three models, and two combined forecasts. It is found that the combination based on posterior probabilities yields accurate forecasts. Section 5 concludes.

## 2. The models

This section presents three functional forms that can be used to model and predict S shaped diffusion processes. These forms are chosen because these are extensively used in practice, as they are not nested, and because they all have just three unknown parameters. Denote the cumulative new cases as *y*_*t*_, for *t* = 1,2,3,…, *T*, and denote *m* as the final cumulative number of cases, and *α* > 0 and *β* > 0 as two unknown parameters.

The first is the logistic curve, which is represented by

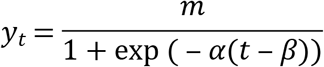

The inflection point, or peak moment of new cases, that is, 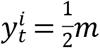 occurs at *t* = *β*. Adding an error term, the parameters (and associated standard errors) can be estimated using Nonlinear Least Squares (NLS). The second is the Gompertz curve, represented by

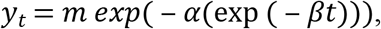

for which 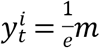, which occurs at 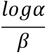. As the inflection point has a cumulative number of cases which is less than 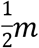, the Gompertz curve is asymmetric. Again, NLS can be used to estimate the parameters. The third and final function is the Bass (1969) growth curve, which is represented by

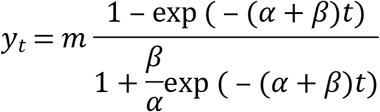

The inflection point is at 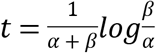 and at the peak, cumulative cases are 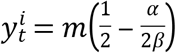. Again, this is an asymmetric curve as the latter expression is smaller than 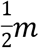. There are various estimation methods for the parameters, but to treat all three models equally, I will consider NLS also for the Bass model.

## 3. Simple Bayesian forecast combination

The three models will be considered for the sample *t* = 1,2,3,…, *T*, and then a one-step-ahead forecast will be created for *T* + 1. Next, the sample moves to *t* = 1,2,3,…, *T* + 1, and a forecast is created for *T* + 2. This will be repeated for *N* forecasts.

Each of the three models gives *N* one-step-ahead forecasts, and these are denoted as 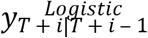, as 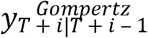 and as 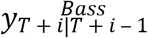, respectively, for *i* = 1,2,3,…,*N*. Additionally, it might be useful to incorporate combined forecasts. A simple average forecast, which, as is found in the literature, is hard to beat in many practical situations, is

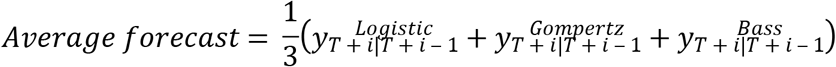

An alternative combination might take into account the quality of the model in the in-sample period. One way to do so is to consider the in-sample posterior probabilities of each of the models given the data. When the prior probability of each of the models is equal, which seems reasonable here, Raftery (1995) shows that the posterior probability (here for the Logistic curve for illustration) can be computed as

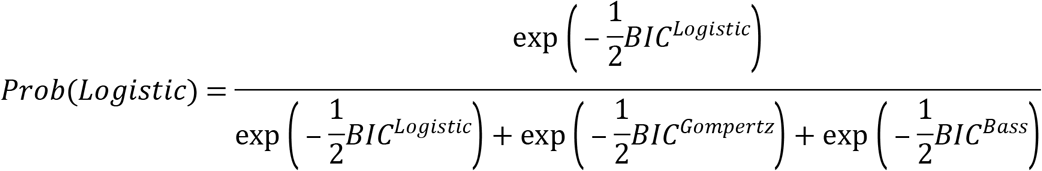

where BIC is the familiar Bayesian Information Criterion (Schwarz, 1978). With these three posterior probabilities, one can construct a Bayesian average forecast simply as

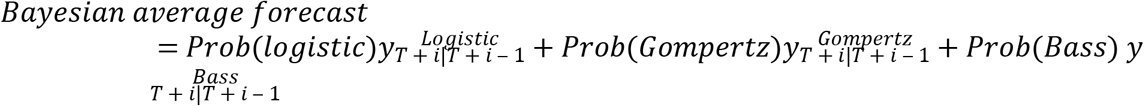

As the forecasts are created recursively, one should also compute for each forecast the BIC again for the recursive samples.

## 4. Covid-19 cases in the Netherlands

To illustrate, consider the data as in Figure 1. The data run from February 27, 2020 (with the first case) to and including May 19, 2020. At first sight, it seems that the inflection point is somewhere in the beginning of April. To have a large enough sample to estimate the parameters, and also to have a large enough sample to evaluate the forecasts, I set *T* at March 31, 2020, which is before the visually suggested inflection point, and hence there are *N* = 49 forecasts to evaluate.

Figure 3 presents the *N* = 49 BIC values, where of course lower values indicate a better fit. Figure 4 gives the three posterior probabilities over time, and it is clear that the posterior probability of the Gompertz model is about twice as large as those for the other two models.

**Figure 3:**
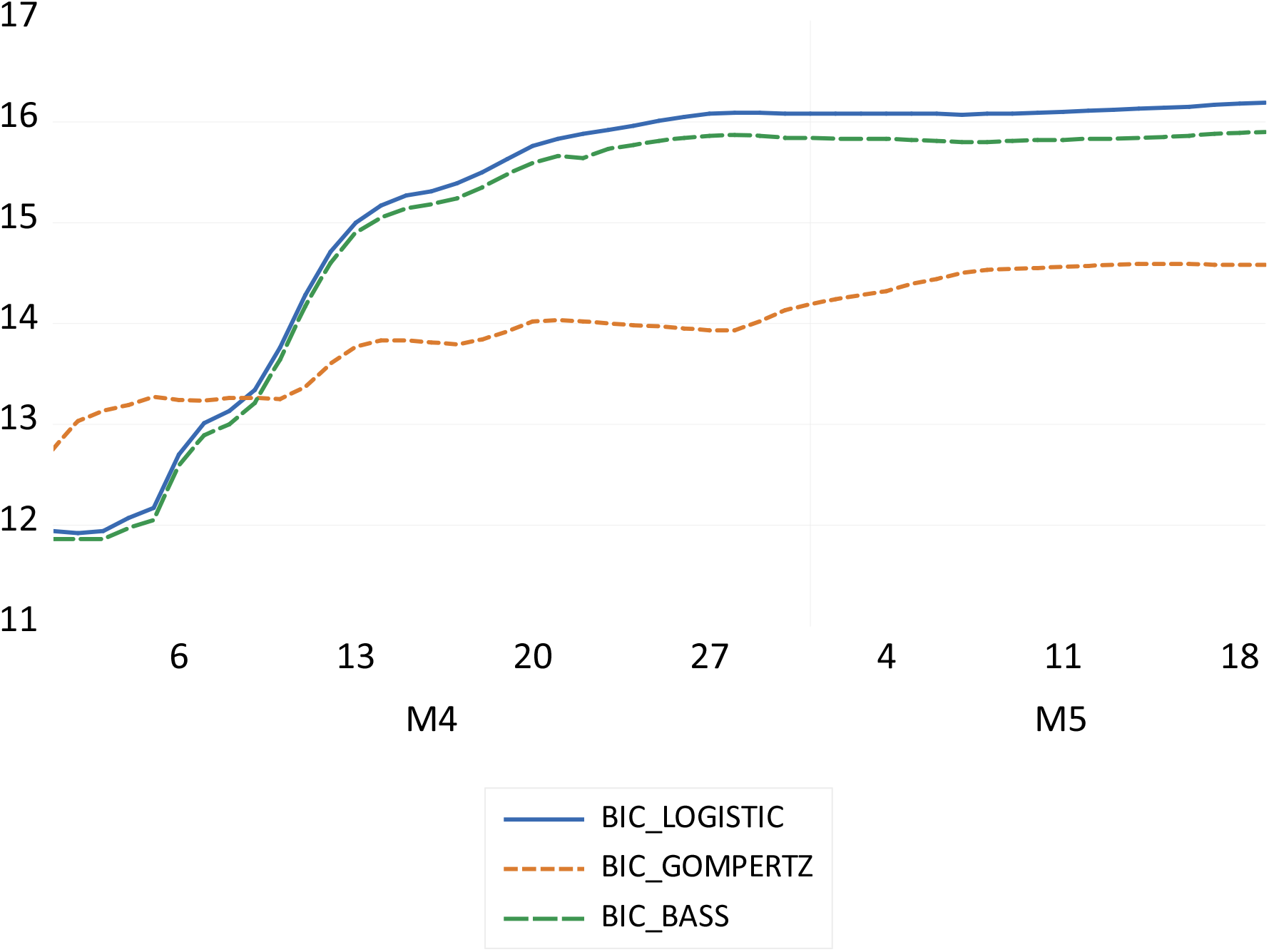
Recursively estimated in-sample BIC values for the three models.

**Figure 4:**
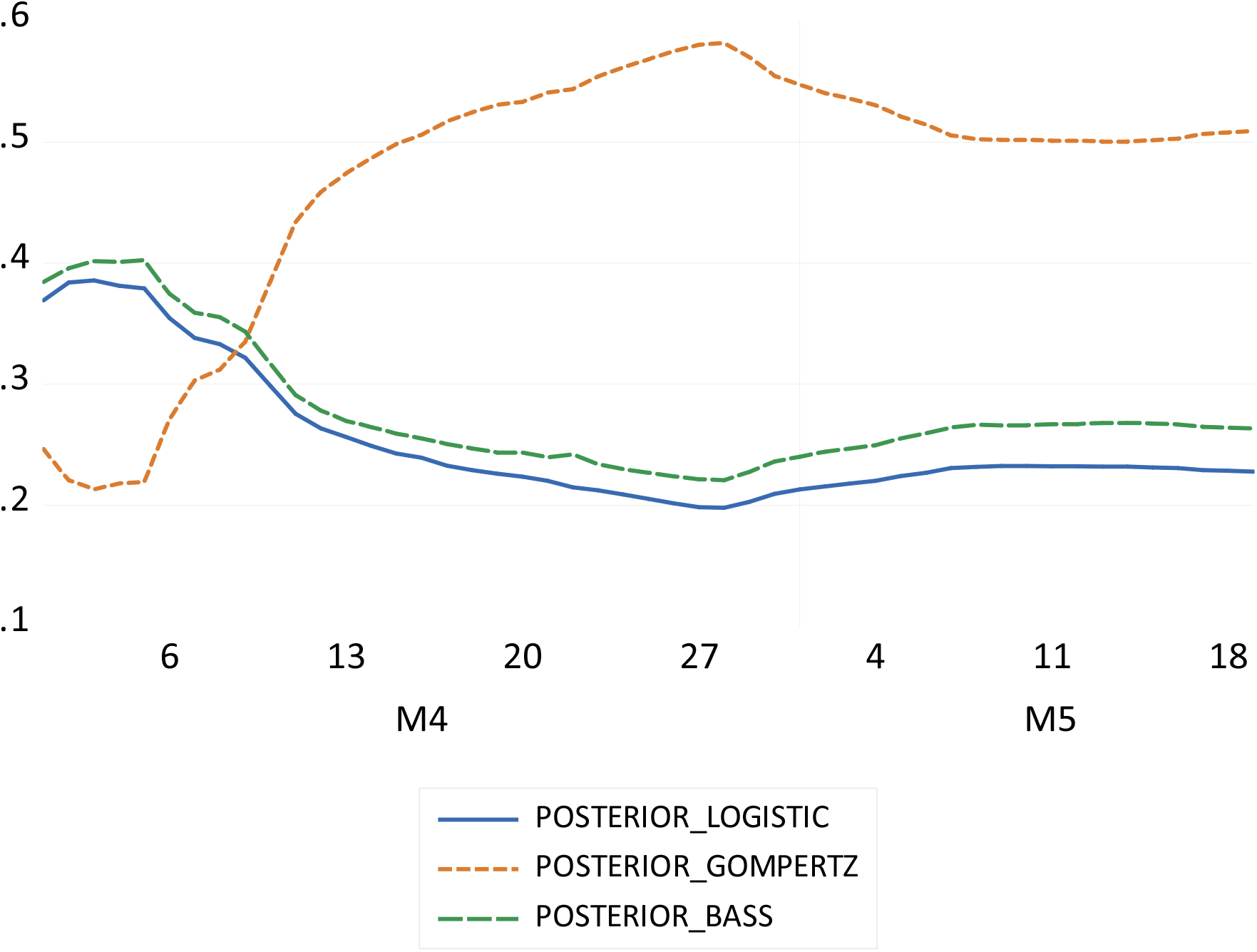
The posterior probabilities for each of the three models, based on the recursively computed BIC values.

Table 1 presents the estimation results for the three models for the full sample with *T* + *N* observations. Looking back from May 19, 2020 it seems that the inflection point occurred somewhere in the first ten days of April 2020.

**Table 1:** Estimation results using NLS (estimated standard errors in parentheses)

| | $m$ | $\alpha$ | $\beta$ | Inflection point at |
| --- | --- | --- | --- | --- |
| Logistic | 44774 (102) | 0.098 (0.001) | 43.33 (0.071) | April 9, 2020 |
| Gompertz | 46324 (195) | 15.08 (0.623) | 0.071 (0.001) | April 4, 2020 |
| Bass | 44838 (104) | 0.001 (0.000) | 0.095 (0.001) | April 9, 2020 |

Table 2 presents the results on forecast accuracy. Clearly, the Logistic and Bass models perform poorly, whereas the Gompertz model gives rather accurate forecasts. The Bayesian average forecast is more accurate than the simple average forecast, and hence it seems useful to incorporate the quality of the in-sample fit in the combined forecast. In terms of absolute forecast errors, the Bayesian combined forecasts are more accurate than the logistic forecasts for 48 of the 49 days, more accurate than the Bass forecasts also in 48 of the 49 days, more accurate than the sample equal weight averages in 41 of the 49 days, but only for 24 of the 49 days more accurate than the Gompertz model forecasts are obtained. This closeness is also reflected by the error measures in Table 2, which indicate that the Gompertz model itself, at least for these data, provides already reasonably accurate forecasts.

**Table 2:** Forecast error measures

|  | Forecast error |  | Absolute forecast error |  |
| --- | --- | --- | --- | --- |
|  | Mean | Median | Mean | Median |
| Logistic | 995 | 998 | 999 | 998 |
| Gompertz | -170 | -325 | 417 | 439 |
| Bass | 874 | 857 | 880 | 857 |
| Average | 567 | 485 | 580 | 485 |
| Bayesian average | 392 | 333 | 435 | 333 |

## 5. Conclusion

This note has implemented a simple algorithm to combine non-nested model forecasts in a Bayesian way. The algorithm only needs the BIC values, and with these, one can compute the posterior probabilities of each of the models. When evaluating forecasts for Covid-19 cases it was found that the Bayesian approach, which takes the in-sample fit of each of the models into account, yielded more accurate forecasts than equal weighted average forecasts.

